# VirusDIP: Virus Data Integration Platform

**DOI:** 10.1101/2020.06.08.139451

**Authors:** Lina Wang, Fengzhen Chen, Xueqin Guo, Lijin You, Xiaoxia Yang, Fan Yang, Tao Yang, Fei Gao, Cong Hua, Yuantong Ding, Jia Cai, Linlin Yang, Wei Huang, Zhicheng Xu, Bo Wan, Jiawei Tong, Chunhua Peng, Yawen Yang, Lei Zhang, Ke Liu, Feiyu Zhou, Minwen Zhang, Cong Tan, Wenjun Zeng, Bo Wang, Xiaofeng Wei

## Abstract

**Motivation:** The Coronavirus Disease 2019 (COVID-19) pandemic poses a huge threat to human public health. Viral sequence data plays an important role in the scientific prevention and control of epidemics. A comprehensive virus database will be vital useful for virus data retrieval and deep analysis. To promote sharing of virus data, several virus databases and related analyzing tools have been created.

**Results:** To facilitate virus research and promote the global sharing of virus data, we present here VirusDIP, a one-stop service platform for archive, integration, access, analysis of virus data. It accepts the submission of viral sequence data from all over the world and currently integrates data resources from the National GeneBank Database (CNGBdb), Global initiative on sharing all influenza data (GISAID), and National Center for Biotechnology Information (NCBI). Moreover, based on the comprehensive data resources, BLAST sequence alignment tool and multi-party security computing tools are deployed for multi-sequence alignment, phylogenetic tree building and global trusted sharing. VirusDIP is gradually establishing cooperation with more databases, and paving the way for the analysis of virus origin and evolution. All public data in VirusDIP are freely available for all researchers worldwide.

**Availability:** https://db.cngb.org/virus/

## 1 Introduction

The COVID-19 pandemic caused by hCoV-19 is a global health emergency and poses a huge threat to human public health (Harapan *et al.*, 2020; Sohrabi *et al.*, 2020; Velavan *et al.*, 2020). Viral sequence data can provide an important scientific basis for the traceability and evolution of viruses (Lu *et al.*, 2020), the development of viral nucleic acid detection reagents, vaccines, and drugs (Lundstrom, 2015; Lundstrom, 2018; Zhang *et al.*, 2020), and play an important role in clinical assistant diagnosis and scientific prevention and control of epidemics.

A one-stop comprehensive virus database will be vital for virus data retrieval and deep analysis. Several organizations at home and abroad have constructed virus databases and developed related analysis tools, such as NCBI Virus (Hatcher *et al.*, 2017), GISAID (Elbe *et al.*, 2017; Shu *et al.*, 2017), COVID-19 Genomics UK Consortium (COG-UK Consortium: https://www.cogconsortium.uk/), China National Center for Bioinformation 2019 Novel Coronavirus Resource (2019nCoVR) (Zhao *et al.*, 2020), Influenza Virus Database (IVDB) (Chang et al., 2007), National Microbiological Science Data Center (NMDC: http://nmdc.cn/). They have made great efforts for archiving, sharing and analysis of virus data. To facilitate virus research and promote the global sharing of virus data, we built VirusDIP at China National GeneBank DataBase (CNGBdb: https://db.cngb.org/) and released it on March 27, 2020.

## 2 Methods

VirusDIP is established and maintained using the Django web framework (https://www.djangoproject.com/), the Python programming language and Pycharm (https://www.jetbrains.com/pycharm/). The Centos7 (https://www.centos.org/) is chosen as the operating system of our server. In terms of system architecture, we use NGINX (http://nginx.org/) to provide static resource access, uWSGI (https://uwsgi-docs.readthedocs.io/en/latest/) to deploy query and download services, Postgresql to store metadata. In terms of data security, the database is deployed in the CNGB (Wang *et al.*, 2019) which has passed the three-level review of information security level protection and the protection capability review of trusted cloud service. Moreover, all services have been deployed with high availability.

## 3 Results

The main functions of VirusDIP includes archive, integration, access, and analysis of virus data (Fig. 1).

**Fig 1.**
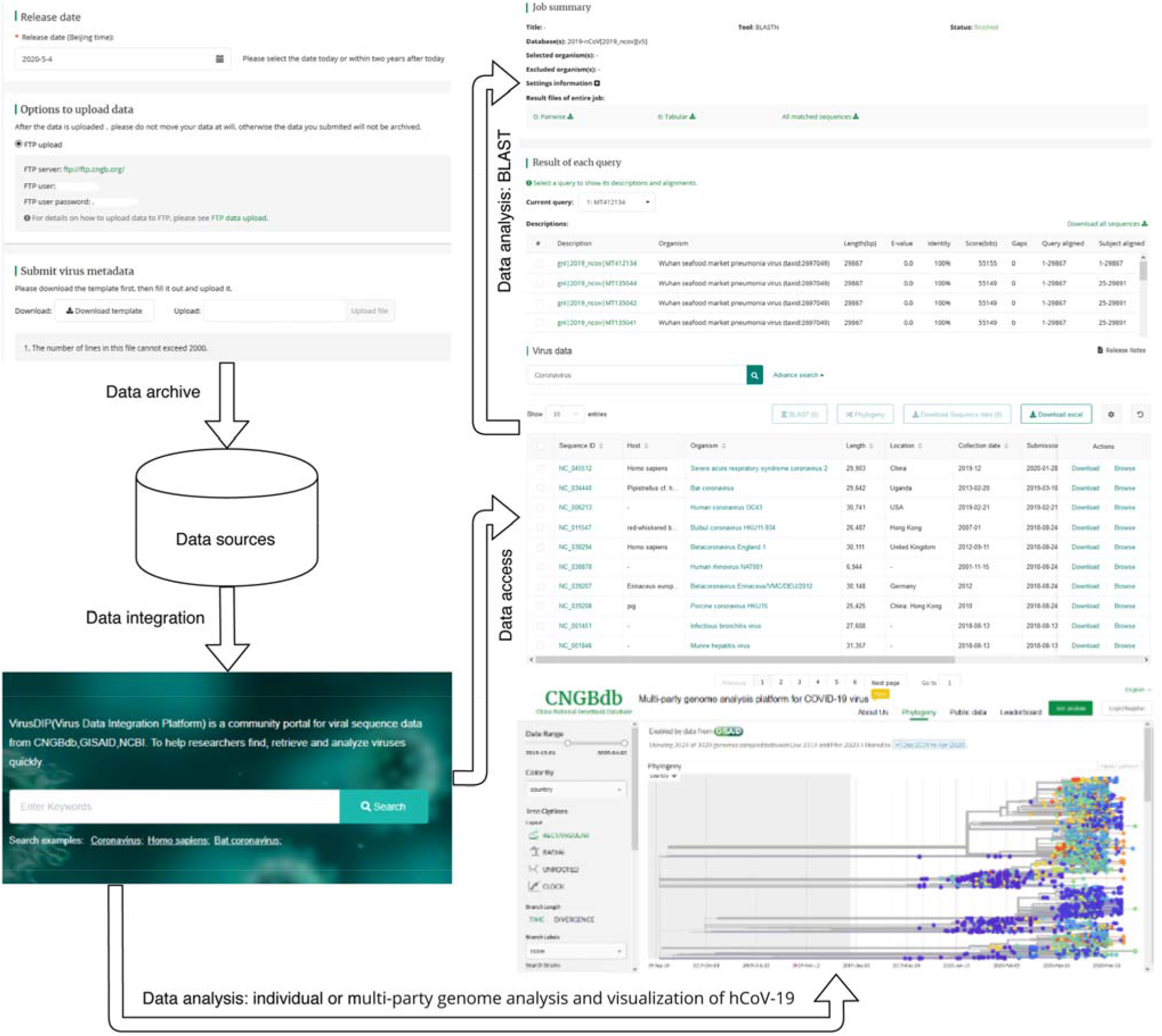
VirusDIP function display. VirusDIP archives virus data submitted by CNGBdb users and integrates these data with public data from different data sources to provide users with free data access. Users can perform BLAST alignment based on the retrieved virus data, and can perform individual and multi-party genome analysis on the hCoV-19 genome data and visually display its phylogeny.

### 3.1 Data archive

We have developed a convenient and rapid submission process for viral sequence data. The submission portal is at https://db.cngb.org/cnsa/init?type=virus. For data compatibility, the virus data standard integrates the virus and pathogen sample data standard of The International Nucleotide Sequence Database Collaboration (INSDC) (Karsch-Mizrachi *et al.*, 2018), the hCoV-19 data standard of GISAID, and the sample data standard of COVID-19 Genomics UK (COG-UK) Consortium. All submitted data must be validated by machine and manual curation to normalize sequences and sample attributes, and ensure data quality.

### 3.2 Data integration and access

VirusDIP currently integrates the virus data submitted by CNGBdb users and shared virus data of GISAID and NCBI. All shared data follows their original data agreements of its source database. On Mar 16, 2020, CNGB and GISAID reached a strategic cooperation and plans to gradually establish a GISAID mirror database in VirusDIP. At present, the hCoV-19 sequence metadata of GISAID can be synchronized to VirusDIP in real time. Table 1 summarizes all integrated data of VirusDIP as of June 4, 2020. At the same time, to make data with the same theme more easily accessible, three datasets based were created, namely hCoV-19 Database (https://db.cngb.org/datamart/disease/DATAdis19/), hCoV-19 potential intermediate host dataset (https://db.cngb.org/datamart/disease/DATAdis21/) and Dataset of bat genome (https://db.cngb.org/datamart/animal/DATAani17/).

**Table 1.** Data sources of VirusDIP

| Data source | Nucleotide sequences | Genomes | Runs |
| --- | --- | --- | --- |
| CNGBdb | - | 75 | 123 |
| GISAID | - | 37,735 | - |
| NCBI | 5,069 | 1 | 7,234 |

All integrated data can be retrieved by entering keywords in the search box on the homepage, such as Coronavirus, Homo sapiens. All data shared by GISAID can be accessed at https://db.cngb.org/gisaid/. Data of hCoV-19 submitted by CNGBdb users or from NCBI can be accessed in COVID-19 (https://db.cngb.org/virus/ncov). The public virus data from NCBI can be accessed in Virus (https://db.cngb.org/virus/virus). Moreover, the table setting function can be used for personalized display and download.

### 3.3 Data analysis

#### 3.3.1 BLAST

VirusDIP has deployed the BLAST sequence alignment tool and established thematic sequence alignment databases for the hCoV-19 and comprehensive virus data to facilitate scientific researchers to quickly perform sequence alignment on a certain virus or other various viruses and analyze the similarity of virus sequences.

#### 3.3.2 Individual and multi-party genome analysis for hCoV-19

Data security and privacy protection are new challenges while promoting the open sharing the biological big data. Blockchain and secure multi-party computation technology have provided a new technical concept for biological big data governance and applied to medical and genomic data analysis (Ahmad *et al.*, 2019; Bogdanov *et al.*, 2018; Vazirani *et al.*, 2019; Veeningen *et al.*, 2018). Based on Nextstrain (Hadfield *et al.*, 2018) and visualization tools, VirusDIP has deployed Multi-party genome analysis platform for hCoV-19, which is the first hCoV-19 analysis tool based on blockchain and secure multi-party computation in China to our knowledge. The tool can display the evolution of the existing hCoV-19 sequences on the platform, including phylogenetic tree and geographic distribution. Users can use their private data and the public data of VirusDIP to perform combined computing to obtain the relative position of private virus sequences on the current public evolution tree, fully guaranteeing data security and promoting the sharing and application of virus data.

## 4 Discussion

VirusDIP has built an efficient batch submission process for virus data and integrated comprehensive virus data from multiple data sources. All integrated virus data can be freely accessed in one stop. VirusDIP also provides users a real-time, dynamic online joint evolution analysis tool for phylogenetic analysis of hCoV-19. VirusDIP is committed to building a comprehensive virus data platform for archive, integration, access, and analysis. In the future, we will continue to improve the data standards for archiving virus data. Moreover, we will gradually develop or deploy new visualization and evolution analysis tools for virus data, such as literature tracking, variation analysis, browser and Phylogeny to further promote the global trusted sharing of virus data and data mining.

In addition, we will continue to strengthen data sharing cooperation with GISAID, expand cooperation with other virus databases and support more projects to promote the establishment of a global cooperation network for life resources.

## Acknowledgements

We gratefully acknowledge all collaborators and authors from originating and submitting laboratories of virus data integrated by VirusDIP.

